# Hybrid peeling for fast and accurate calling, phasing, and imputation with sequence data of any coverage in pedigrees

**DOI:** 10.1101/228999

**Authors:** Andrew Whalen, Roger Ros-Freixedes, David L Wilson, Gregor Gorjanc, John M Hickey

## Abstract

In this paper we extend multi-locus iterative peeling to be a computationally efficient method for calling, phasing, and imputing sequence data of any coverage in small or large pedigrees. Our method, called hybrid peeling, uses multi-locus iterative peeling to estimate shared chromosome segments between parents and their offspring, and then uses single-locus iterative peeling to aggregate genomic information across multiple generations. Using a synthetic dataset, we first analysed the performance of hybrid peeling for calling and phasing alleles in disconnected families, families which contained only a focal individual and its parents and grandparents. Second, we analysed the performance of hybrid peeling for calling and phasing alleles in the context of the full pedigree. Third, we analysed the performance of hybrid peeling for imputing whole genome sequence data to the remaining individuals in the population. We found that hybrid peeling substantially increase the number of genotypes that were called and phased by leveraging sequence information on related individuals. The calling rate and accuracy increased when the full pedigree was used compared to a reduced pedigree of just parents and grandparents. Finally, hybrid peeling accurately imputed whole genome sequence information to non-sequenced individuals. We believe that this algorithm will enable the generation of low cost and high accuracy whole genome sequence data in many pedigreed populations. We are making this algorithm available as a standalone program called AlphaPeel.

## Introduction

In this paper we extend multi-locus iterative peeling to be a computationally efficient method for calling, phasing, and imputing low coverage sequence data in large pedigrees. In the past few years the use of genomic data has expanded greatly. The widespread genotyping of animals empowers breeding via genomic selection (Meuwissen et al., 2001, 2016) and biological discovery via genome wide association studies (Burton et al., 2007; Visscher et al., 2017). The accuracy of genomic selection and the power of genome wide association studies depend on both the number of individuals that have genomic data and its density (e.g., Daetwyler et al., 2008; Hayes et al., 2009; Hickey et al., 2014; Gorjanc et al., 2015). The goal is then to generate genomic data on as many individuals as possible at as high of a density as possible with the upper limit being the presence of whole genome sequence on hundreds of thousands or millions of individuals (Hickey, 2013; Daetwyler et al., 2014; Veerkamp et al., 2016).

Even though the cost of obtaining whole genome sequence data on an individual has decreased, it is still prohibitively expensive to obtain high coverage whole genome sequence data on tens of thousands of individuals. An emerging strategy in breeding populations is to obtain a mix of high and low coverage sequence data on a subset of individuals, and then propagate that information between related individuals to call whole genome sequence genotypes for all population members, some of which may only have SNP array genotype data (Hickey, 2013). This strategy exploits the high degree of relatedness and thus haplotype sharing between individuals in a breeding population, meaning that a haplotype can be inferred at high accuracy by low coverage sequencing of different individuals that share the haplotype. Algorithms have already been developed for selecting the individuals to sequence in such a context (Cheung et al., 2014; Gonen et al., 2017; Ros-Freixedes et al., 2017). What remains to be developed is a method for efficiently propagating the information from sequence data between related individuals.

Past methods for using mixed coverage sequence data to call, phase, and impute genotypes have primarily exploited linkage disequilibrium, e.g. MaCH (Li et al., 2010), Beagle (Browning and Browning, 2016, 2007). Linkage disequilibrium based methods perform well, particularly in human settings where individuals are mostly unrelated and there is limited pedigree data. However, these methods do not exploit the large amount of information available when pedigrees are available (but see, Browning and Browning, 2009; O’Connell et al., 2014). In contrast, pedigree based methods can have a higher accuracy and lower computational cost than linkage disequilibrium based methods, particularly in populations with closely related individuals and accurate pedigrees across multiple generations (e.g., Hickey et al., 2011; Cheung et al., 2013; VanRaden et al., 2015). Pedigree based methods are particularly appealing for mixed coverage sequence data on relatives, due to being able to collapse information across the long haplotype segments shared between individuals, their ancestors and their descendants.

Single-locus and multi-locus peeling are two pedigree-based methods that model an individual’s haplotype based on the haplotypes of their parents and offspring. There is a large body of literature on peeling methods in genetics (e.g., Elston and Stewart, 1971; Cannings et al., 1976, 1978; Lander and Green, 1987; Fernández et al., 2001; Totir et al., 2009; Cheung et al., 2013) and related methods in other areas (e.g., Lauritzen and Sheehan, 2003; Bishop, 2007; Koller and Friedman, 2009). Since our interest is in efficient methods that could handle whole genome sequence data in multi-generational pedigrees with loops, we focus on approximate (iterative) peeling methods, in particular to the single-locus method of Kerr and Kinghorn (1996) and multilocus method of Meuwissen and Goddard (2010). In single-locus peeling all loci are treated independently and so linkage between loci is not exploited. In contrast multi-locus peeling tracks the linkage between loci allowing for information at one locus to be used at a neighbouring locus, which has a large potential with sequence data. Although multi-locus peeling is exploiting more information and is therefore more accurate, it is computationally more expensive due the high cost of calculating the segregation estimates at each locus, and currently is ill-suited for whole genome sequence data.

In this paper we present a hybrid peeling method that is scalable to whole genome sequence data on tens of thousands of individuals. In hybrid peeling segregation estimates are calculated on a small subset of loci, and then fast single-locus style peeling operations are used on the remaining loci. This approach exploits the benefits of using linkage from multi-locus peeling while still being able to scale to whole genome sequence data on thousands of animals. In what follows we first present the hybrid peeling method, and then present results of its performance on a synthetic dataset based on a real commercial pig population with 60,000 animals on a single chromosome with 700,000 segregating loci. We found that hybrid peeling substantially increases the number of genotypes that were called and phased by leveraging sequence information on related individuals. The calling rate and accuracy increased when the full pedigree was used compared to a reduced pedigree of just parents and grandparents. Finally, we found that hybrid peeling accurately imputes whole genome sequence information to non-sequenced individuals. We are making this algorithm available as a standalone program called AlphaPeel.

## Materials and Methods

### Peeling methods

Peeling is a method for inferring the genotype and phased alleles of an individual based on their own genotype information and the genotype information of their ancestors and descendants. The genotype information can be partially or fully (in)complete or even incorrect for some pedigree members. This inference problem is computationally intractable when considering whole genome sequence in the context of large multi-generational pedigrees with loops (Cannings et al., 1978; Lauritzen and Sheehan, 2003; Piccolboni and Gusfield, 2003; Totir et al., 2009). Iterative peeling approximates this problem through a series of peeling up and peeling down operations (Van Arendonk et al., 1989; Kerr and Kinghorn, 1996; Meuwissen and Goddard, 2010). In the following we refer to iterative peeling simply as peeling. In a peeling up operation information from an individual’s descendants and their mates is used to infer the individual’s alleles. In a peeling down operation information from an individual’s ancestors is used to infer the individual’s alleles. Repeated peeling operations propagates genetic information between distant members of a pedigree.

Peeling relies on a model of how alleles are transmitted between a parent and their offspring. Single-locus and multi-locus peeling differ in how they model the transmission of alleles. In single-locus peeling, both parental alleles are assumed to be inherited with equal probability at all loci. In multi-locus peeling, it is assumed that there is a high probability that the nearby loci are inherited from the same paternal gamete. To enable the sharing of information between loci, multi-locus peeling estimates the segregation at each locus, the likelihood that each pair of grandparental gametes was inherited at a locus. Hybrid peeling is a computationally efficient approximation to multi-locus peeling. Like multi-locus peeling it utilizes information from nearby loci to determine which allele is inherited at a locus. Unlike multi-locus peeling, it only estimates segregation on a small subset of loci, and linearly interpolates segregation estimates at un-evaluated loci.

We describe these peeling operations in detail below. For single-locus peeling we follow the previous work of Kerr and Kinghorn (1996) and for multi-locus peeling we follow the previous work of Meuwissen and Goddard (2010).

### Single-locus peeling

In single-locus peeling we estimate the likelihood of each of an individual’s alleles at a locus as the product of their parents’ alleles (anterior), offsprings’ alleles (posterior), and their own genomic data (penetrance). For a biallelic loci, we have a set of four possible ordered pairs of alleles (aa, aA, Aa, AA), where the first allele in each pair is inherited from the father and the second allele is inherited from the mother. The probability that individual *i* has alleles *h_i_* is:

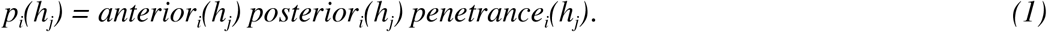

We examine each of these terms separately.

The penetrance term gives the likelihood that an individual has a given set of alleles based on the available genomic data, obtained either from a SNP array or sequencing. If no information is available, we set the penetrance to a constant value, i.e., *penetrance_i_(h_j_) = 1*. If we have SNP array data, we set *penetrance_i_(h_j_) = 1-ε* if *h_i_* is consistent with the genotype on the SNP array, and *penetrance_i_(h_j_) = ε* otherwise, where *ε* accounts for a small error rate in SNP array genotype data. If we have sequencing data with n_ref_ sequence reads of the reference allele, a, and n_alt_ sequence reads of the alternative allele, A, then:

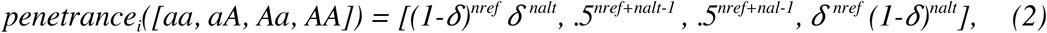

where *δ* accounts for a small error rate in sequence data.

The anterior estimate captures the information about an individual’s haplotypes gained from their parents’ haplotypes. If an individual does not have any genotyped parents, then we use the minor allele frequency, *p*, to calculate the anterior estimate:

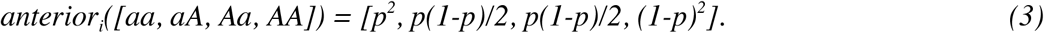

For an individual with parents the anterior estimate is:

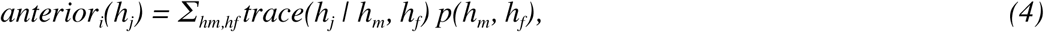

where *p(h_m_*, *h_f_)* is the joint probability that the mother has alleles *h_m_* and the father has alleles *h_f_*, The trace is a function that gives the likelihood that the child inherits alleles *h_i_* given their parent’s alleles, i.e., *trace(h_j_* | *h_m_*, *h_f_)* = *p(h_j_* | *h_m_*, *h)*. Examples of the trace function when inheriting from a single parent are given in Table 1(a). The joint probability of the parental alleles is calculated by combining the anterior and posterior estimates for both parents except for the information that pertains to individual *i*. This gives:

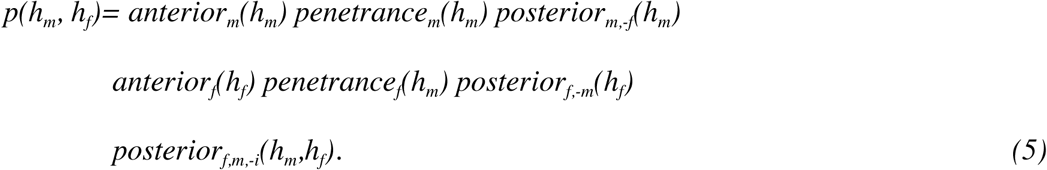

**Table 1.** Examples of the trace function under single-locus peeling (a) and multi-locus peeling (b) when the child inherits the grandpaternal (first) allele.

| (a) Equal likelihood of inheritance |  |  | (b) Grandpaternal inheritance |  |  |
| --- | --- | --- | --- | --- | --- |
| Parental haplotype | Inherited allele | Trace probability | Parental haplotype | Inherited allele | Trace probability |
| aa | a | 1 | **aa** | a | 1 |
| aA | a | 0.5 | **aA** | a | 1 |
| Aa | a | 0.5 | **Aa** | a | 0 |
| AA | a | 0 | **AA** | a | 0 |
| aa | A | 0 | **aa** | A | 0 |
| aA | A | 0.5 | **aA** | A | 0 |
| Aa | A | 0.5 | **Aa** | A | 1 |
| AA | A | 1 | **AA** | A | 1 |

The first line calculates the probability of the mother’s alleles, *h_m_*, independent of shared children with *f*. The second line calculates the probability of the father’s alleles, *h_f_*, independent of shared children with m. The third line calculates the probability of both parents’ alleles based on their shared children except for individual *i*.

There are two types of posterior terms. First, *posterior_m,f_*, is the joint probability of two parents’ alleles, *m* and *f*, based on all their shared children. Second, *posterior_m_* is the probability of a single parent’s alleles based on all their mates and children. We can calculate *posterior_m,f_* by:

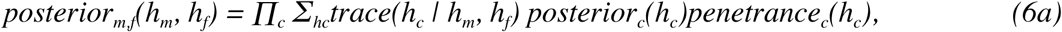

which is the product of the probability that a child, *c*, inherits alleles *h_c_*, based on their parent’s alleles, marginalized over the possible alleles for c, and multiplied across all children. We can then calculate *posterior_m_(h_m_)* as the product of the *posterior_m,f_(h_m_*,*h_f_)* for all of the mates of *m* marginalized over the likelihood that *k* has alleles *h_k_*:

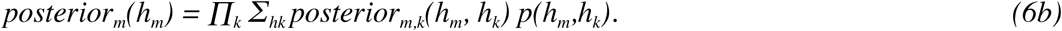

The remaining terms are calculated by removing the individuals that relate to them in the equations:

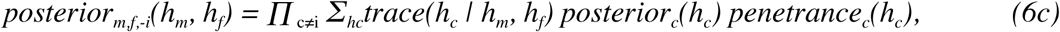

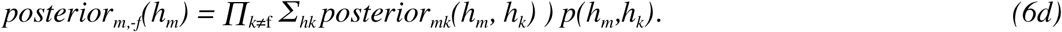

Together the posterior, anterior, and penetrance terms give the probability of individual’s alleles (Equation 1). Information from siblings, parents, and grandparents is contained in the anterior term. Information from children, grandchildren, and their mates is contained in the posterior term. An individual’s own information is only counted a single time, in the penetrance function. When estimating the genotype of a set of parents in the anterior term, the focal individual’s penetrance and anterior terms are excluded from the calculation (Equation 5), which ensures that information from an individual is included in only the anterior or posterior term but not both.

To perform peeling we initialize the population by setting all the posterior terms to a constant value, i.e. 1. We first peel down, updating the anterior terms for all individuals. We then peel up the pedigree, updating the posterior terms for all individuals. These peeling operations are repeated until the allele estimates for all of the individuals in the population converge. There are two model parameters that need to be estimated, the minor allele frequency, *p*, and error rates, *ε* and *δ*. We found that an easy way to update them is by setting them equal to their observed values after each pair of peeling (up and down) operations. We tested using a single error rate for all loci or using a locus specific error rate and found that the locus specific error rate lead to a slight increase in accuracy and so used a locus specific error rate for *ε* and *δ*. Due to the dependence of the anterior terms and posterior terms on the anterior terms and posterior terms of other individuals in the population, the order in which they are updated is important and can decrease the overall number of peeling operations that need to be performed. We follow the updating pattern given in Kerr and Kinghorn (1996) by first updating the anterior terms for the oldest individuals in the population, and then updating the anterior terms for their children and their children’s children. The posterior estimates are updated in reverse order; from the most recent generation to the most distant.

### Multi-locus peeling

Multi-locus peeling extends single-locus peeling by modifying the trace function to be sensitive to which grandparental gamete was likely to have been inherited at nearby loci. In single-locus peeling we assume that each parental allele is inherited with equal probability, and that the alleles at neighbouring loci are inherited independently. This is not the case; due to the small number of recombinations per chromosome, children inherit grandparental gametes in large blocks from their parents. This means that if we know which grandpaternal gamete a child inherits at one locus, we can also know which gamete they likely inherit from at nearby loci. In the context of the peeling operations, if we know which grandpaternal gamete a child is inheriting from, we can update the peeling operations so that only the alleles from that gamete will be transmitted, as in Table 1b. Uncertainty in haplotype inheritance can be incorporated in the model by marginalizing over possible inherited gametes.

More formally, we track the set of inherited haplotypes in terms of a segregation estimate, which gives the likelihood that a child inherits the each of the four possible pairs of grandpaternal gametes (pp, pm, mp, mm); relating to whether the father (first allele) or the mother (second allele) passes their grandpaternal (p) or grandmaternal (m) gamete at that locus. We can then build the trace function by marginalizing over segregation estimates:

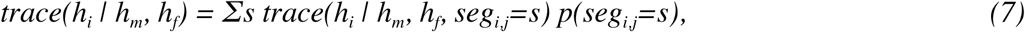

where *p(seg_ij_*,=*s)* is the likelihood that individual *i* has segregation *s* at locus *j. trace(h_i_* | *h_m_*, *h_f_*, *seg_ij_*=*s)* is the likelihood that the child inherits allele *h_i_* given their parental allele and their segregation (see Table 1b for an example). To perform peeling, we substitute the trace function in Equations 4, 6a-d with the trace function Equation 7.

The segregation estimate at each locus is calculated by measuring how well the segregation models the current locus and how well the segregation estimate matches that of adjacent loci:

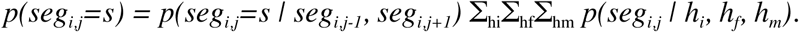

The first term accounts for the recombination rate between loci and the second term accounts for the additional information gained from the genotype estimate at the current allele:

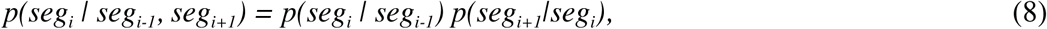

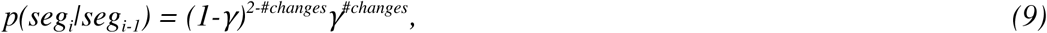

where, *#changes* is the number of gametes that switch (up to 2) between *seg_i_* and *seg_i-1_*, and *γ* is recombination rate. We estimate *p(seg_i_*|*seg_i-1_*, *seg_i+1_)* using the forward-backward algorithm (Rabiner, 1989). To calculate the likelihood of a segregation estimate given the observed data at a locus, we marginalize over possible allele combinations:

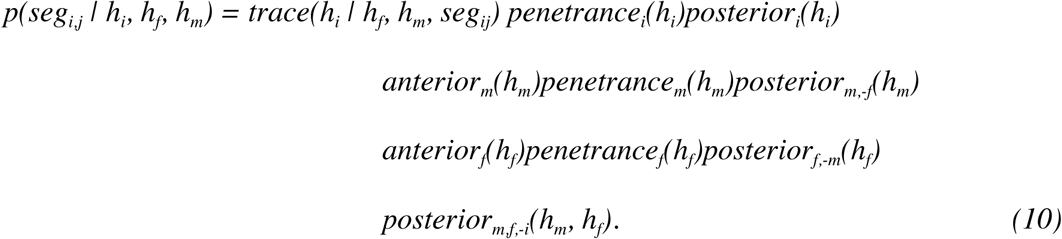

The first line is the likelihood of the child’s alleles, the second is the likelihood of the mother’s alleles, the third is the likelihood of the father’s alleles, and the fourth is the joint likelihood of the parents’ alleles.

This algorithm is performed in a series of forward-backward passes where at each locus all individuals in the population are updated by the peeling up and peeling down operation. Segregation estimates are then re-estimated for each individual. At the end of each pass we updated the recombination rate, *γ*, error rate, *ε* and *δ*, and minor allele frequency, *p*, by setting them to their observed values. Similar to the error rate we found that using a locus specific recombination rate slightly increased accuracy and so used a locus specific recombination rate. We found that between 10-20 cycles was enough to obtain convergence in large multi-generational livestock pedigrees with 60,000+ members.

### Hybrid peeling

Hybrid peeling is a computationally efficient approximation to multi-locus peeling. In preliminary work we found that the primary computational cost of multi-locus peeling stemmed from updating the segregation estimates, Equation 10. When evaluating many loci on a chromosome we should expect that the segregation estimates at nearby loci should be identical. Because of this, it should be possible to evaluate the segregation estimates at only a subset of loci, and interpolate segregation estimates on the remaining loci. These estimates can then be used to create a new trace function for peeling operations.

More formally, we divide the set of loci into two sets, A and B, with |A| ≪ |B|, e.g., A is a subset of loci on a high-density SNP array, and B is the entire set of segregating loci. We perform multi-locus peeling on A to calculate segregation estimates. We then perform single-locus peeling on B using Equation 7 as the trace function with interpolated segregation estimates:

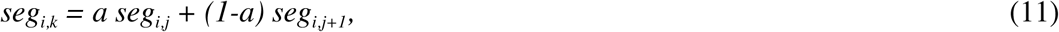

where *j* and *j+1* are the loci in the set A that flank locus *k*, and *a* is the proportional distance between locus *k* and locus *j*:

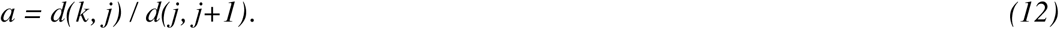

Distance can be calculated either in terms of base pairs, centiMorgans, or number of intermediary loci. The exact measure of distance should only have a minimal impact on performance: if a sufficiently large number of loci is used in the set A then adjacent segregation estimates should be nearly equal, i.e., seg_i,j_ = seg_i,j+1_, leading Equation 11 to reduce to *seg_i,j_* and no longer depend on the distance metric used.

The aim of the hybrid technique is to make multi-locus peeling more computationally tractable when applying it to large pedigrees. We evaluate the performance of this algorithm on a synthetic dataset.

### Analysis

We examined the performance of hybrid peeling for calling, phasing, and imputing alleles with sequence data of different coverages in pedigrees. To perform these analyses, we simulated genomes for 64,598 animals using a multi-generational pedigree derived from a real commercial pig breeding line. We assumed some animals had high-density or low-density SNP array genotypes from routine genomic selection. In addition, we generated mixed coverage sequence data for a subset of focal animals. We then carried out three sets of analyses. First, we analysed the performance of hybrid peeling in calling and phasing in disconnected families, families which contained only a focal animal and its parents and grandparents. Second, we analysed the performance of hybrid peeling in calling and phasing in the context of the full pedigree. Third, we analysed the performance of hybrid peeling for whole genome sequence imputation. In the following we describe in detail how we simulated and analysed data.

### Data

Genomes were generated using the Markovian Coalescent Simulator (MaCS) (Chen et al., 2009) and AlphaSim (Faux et al., 2016). We generated 1,000 base haplotypes for each of 10 chromosomes, assuming a chromosome length of 10^8^ base pairs, a per site mutation rate of 2.5× 10^−8^, a per site recombination rate of 1 × 10^8^, and an effective population size (N_e_) that varied over time in accordance with estimates for a livestock population (MacLeod et al., 2013). The resulting haplotypes had about 700,000 segregating loci per chromosome. On each of the chromosomes we designated 2,000 evenly distributed loci as markers on a high-density SNP array and a subset of 500 as markers on a low-density SNP array.

We used AlphaSim to drop the base haplotypes through a multi-generational pedigree of 64,598 animals from a real commercial pig breeding line. We assigned SNP array data to animals in line with routine genotyping for genomic selection in the population; 45,592 animals were genotyped with high-density SNP array, 11,015 animals were genotyped with low-density SNP array, and 7,991 animals were not genotyped. We generated sequence data in line with the strategies implemented in the population. The goal was to use roughly $300,000 worth of resources to sequence and impute the entire population. First, the top 475 sires (all sires with more than 25 progeny) were sequenced at 2x. Second, AlphaSeqOpt (Gonen et al., 2017) was used to identify focal animals and their parents and grandparents (211 in total) to sequence and the coverages they should be sequenced at. AlphaSeqOpt was run using the high-density SNP array data on all chromosomes with an option to assign an individual sequencing coverage of either 1x, 2x, 15x, or 30x, and a total budget of $71,000. Third, the top 50 dams (based on number of offspring and grandoffspring with and without a sequenced sire) were sequenced at 2x and the next 450 dams were sequenced at 1x. Finally, AlphaSeqOpt2 (Ros-Freixedes et al.) was used to identify 800 individuals to be sequenced at 1x, to top-up the accumulated coverage of common haplotypes to 10x. In total, we generated sequenced data for 1,912 animals at a range of coverages for a cost of $289,312. We partitioned this data into three sequencing sets: i) the *focal* identified with AlphaSeqOpt, ii) the *focal plus low coverage sires* which also included the top 475 sires, and iii) *focal plus all low coverage individuals* which included all the sequenced animals. A breakdown of the total cost and sequencing coverage by these sets is given in Table 2. We assumed that the cost of obtaining a DNA library for an individual was $39 and the cost of sequencing that library for an individual at 1x was $68, at 2x was $136, at 15x was $408, and at 30x was $816. The costs were assumed to be non-linear to reflect current industry costs.

**Table 2.**
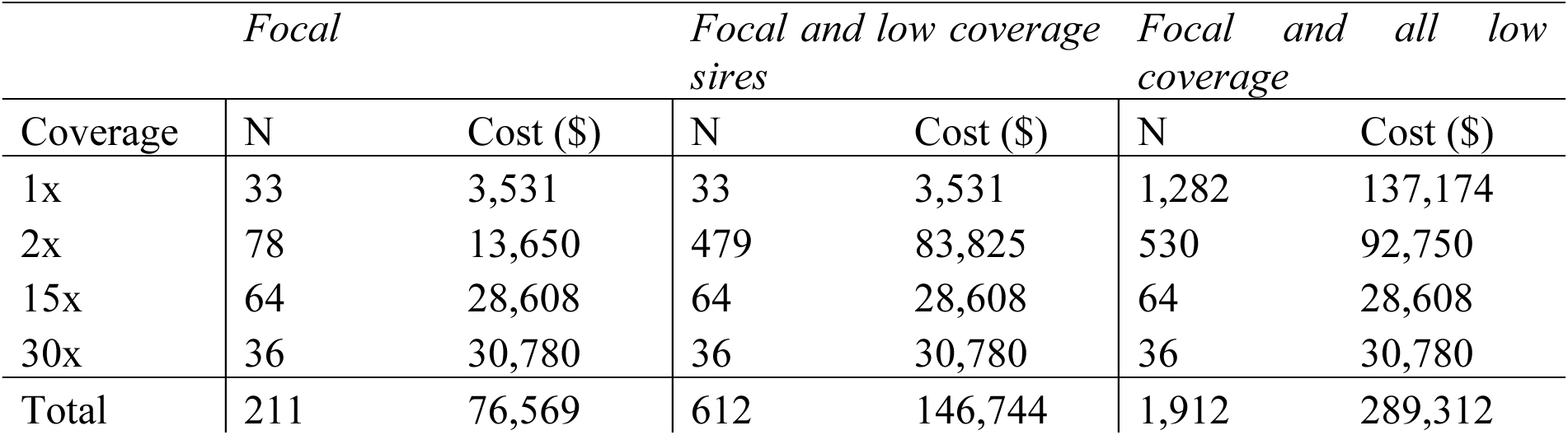
Number of sequenced animals and cost by sequence coverage and the three sequencing sets.

Sequence data was simulated by sampling sequencing reads for the 700,000 segregating loci on the chromosome 10. The number of reads was generated using a Poisson-Gamma distribution which allowed the number of sequence reads per locus to vary along the genome and between individuals (Li et al., 2010). First, a sequenceability (*γ_j_*) of each of the 700,000 loci along the genome was sampled from a gamma distribution, with shape and scale parameters respectively equal to α =4 and 1/α = .25. Second, the number of reads (*r*_i,j_) per individual *i* at locus *j* was then sampled from a Poisson distribution with mean equal to μ_i,j_=X_i_γ_j_, where *x*_i_ was the targeted coverage for individual. Third, sequencing reads were generated by randomly sampling alleles from the two gametes of individual *i* at locus *j*, accounting for a sequencing error (ε = 0.001).

### Calling and phasing in disconnected families

We tested the ability of hybrid peeling to call genotypes and phase alleles in sequenced individuals using information from their parents and grandparents. For this we selected 10 disconnected families (consisting of a focal individual and its parents and grandparents) from the full pedigree, and analysed the effect of sequencing coverage on our ability to call and phase the individual’s genotypes. To perform this, we ran the hybrid peeling when the focal individual was sequenced at 1x, 2x, 5x, 15x, or 30x coverage, and when its parents or grandparents were sequenced at 0x, 1x, 2x, 5x, 15x, or 30x coverage. We generated data for each of these scenarios separately. We assumed that all of the parents or all of the grandparents were sequenced at the same coverage, and that all family members had high-density SNP array data.

To call genotypes and phased alleles, we extracted the allele probabilities generated by hybrid peeling and made a call if the probability of an allele was greater than a pre-defined threshold. For all analyses we used a calling threshold of .98. Scenarios were compared on the percentage of called genotypes (genotype yield) and phased alleles (phase yield).

### Calling and phasing with the full pedigree

Next, we tested the ability of hybrid peeling to call genotypes and phase alleles in sequenced individuals using information from the full pedigree. To perform this, we ran hybrid peeling twice. First, we ran it separately for each disconnected family, consisting of an individual, their parents, and their grandparents, with (potentially missing or low coverage) SNP array and sequence data. Second, we ran it with SNP array and sequence data on all individuals in the pedigree. The sequencing coverage for each individual was determined by their coverage in the *focal and all low coverage* condition. We compared the genotype and phase yield between runs and compared the correlation between individual’s called genotypes and the true genotypes (genotype accuracy) and correlation between individual’s phased alleles and the true phase/haplotype (phase accuracy) between runs.

### Imputing whole genome sequence

Last, we tested the ability of hybrid peeling to impute whole genome sequence for non-sequenced individuals in the full pedigree. We ran hybrid peeling on all of the individuals in the full pedigree, using all available sequence and SNP array data. Hybrid peeling was run three times, using either the sequence coverages from the *focal, focal and low coverage sires*, or *focal and all low coverage* conditions. Imputation accuracy was measured as correlation between an individual’s imputed dosages and the true genotypes.

## Results

Overall, we found that hybrid peeling had high yield and accuracy for called genotypes and phased alleles. It also had a high accuracy of imputing whole genome sequence data to non-sequenced individuals.

### Calling and phasing in disconnected families

We found that hybrid peeling gave high yield and accuracy of called genotypes and phased alleles even in the presence of low coverage sequence reads. The results of these simulations are given in Figure 1.

**Figure 1.**
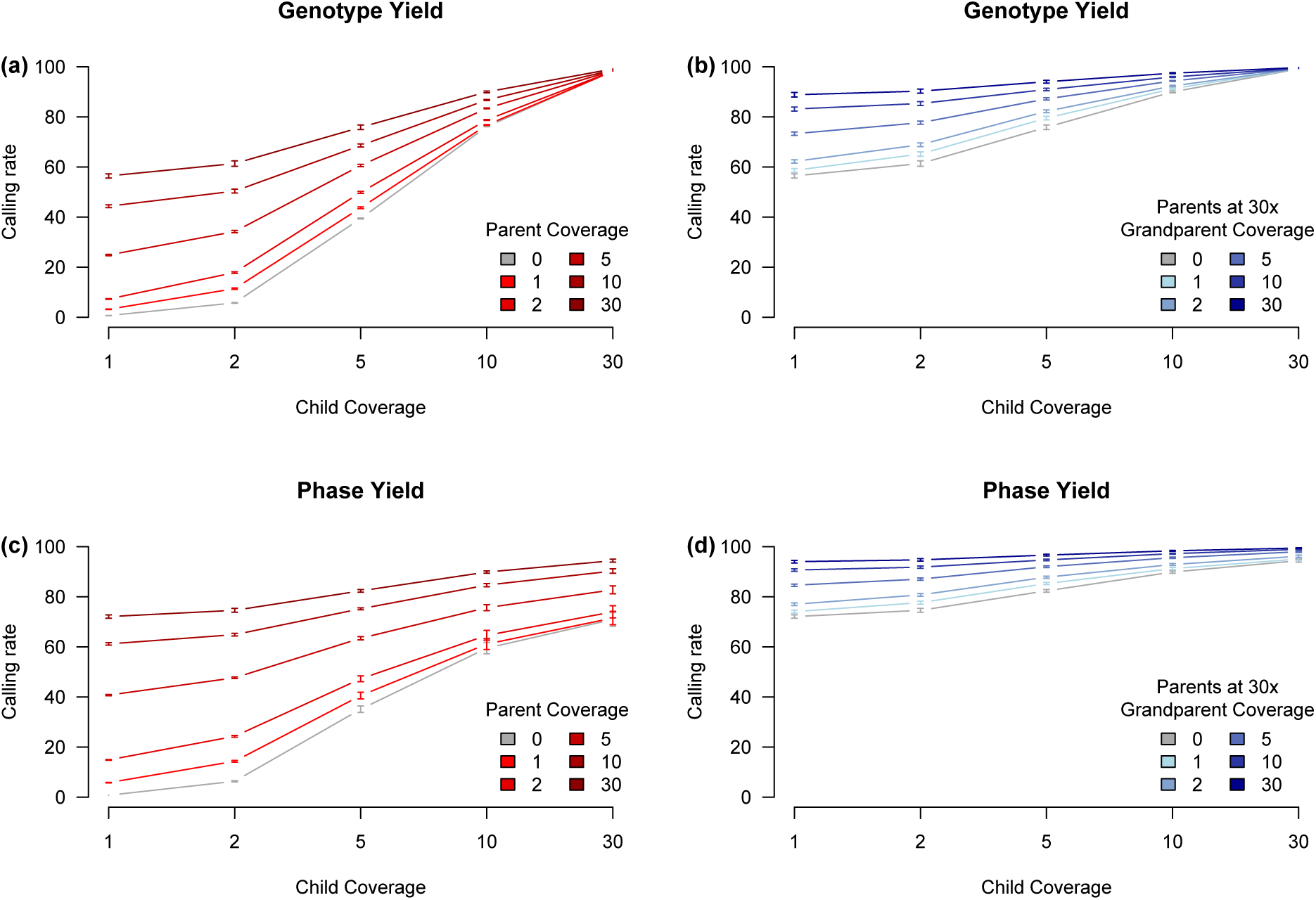
Genotype and phase yield while varying coverage in the focal individual and its parents and grandparents. Panels (a) and (b) give the percentage of called genotypes while varying (a) the coverage in parents and (b) the coverage in grandparents. Panels (c) and (d) give the percentage of phased alleles while varying (c) the coverage in parents and (d) the coverage in grandparents. In panels (b) and (d) the coverage in parents was constant at 30x. In all four panels the accuracy was > .98. Error bars represent plus or minus one standard error based on ten replications.

The primary determiner of genotype yield was the individual’s own degree of sequencing coverage. If neither the individual’s parents nor grandparents were sequenced, then if the individual was sequenced at 1x the genotype yield was 0.6%, and increased to 5% at 2x, 39% at 5x, 76% at 10x, and 98% at 30x. These values greatly increased if the parents were sequenced at high coverage. If the individual’s parents were both sequenced at 30x, then the genotype yield was 56% at 1x, 61% at 2, 75% at 5x, 90% at 10x, and 99% at 30x. Adding in additional coverage on grandparents increased accuracy even if the parents had 30x coverage. If both the parents and the grandparents had 30x coverage then the genotype yield was 88% at 1x, 90% at 2x, 94% at 5x, 97% at 10x, and 99% at 30x. In all cases, the ratio of correctly called genotypes to incorrectly called genotypes was greater than .995 (median .999).

A similar pattern of results was found when evaluating phase yield. In this case, although an individual’s own sequencing coverage was an important determiner for phase yield, high coverage on both the parents and the grandparents were needed to phase all the alleles. If neither the individual’s parents nor grandparents were sequenced, then the phase yield was .7% at 1x, 6% at 2x, 35% at 5x, 59% at 10x, and 70% at 30x. The low phase yield at 30x is due to the inability to phase heterozygous loci without information from relatives. Sequencing the parents at high coverage substantially increased the phase yield, and continued to do so even if the individual was sequenced at high coverage. If the parents of the individual were sequenced at 30x, then the phase yield was 72% at 1x, 74% at 2x, 82% at 5x, 89% at 10x and 94% at 30x. If both the individual’s parents and grandparents were sequenced at 30x, then the phase yield increased to 94% at 1x, 95% at 2x, 96% at 5x, 98% at 10x, and 99% and 30x. In all cases, the ratio of correctly phased alleles to incorrectly phased alleles was greater than 0.989 (median .999).

### Calling and phasing with the full pedigree

We examined the effect of using all sequence data and the full pedigree on calling genotype and phase yield and accuracy of sequenced individuals. The gains in yield and accuracy in comparison to using data from disconnected families are plotted in Figure 2. We found that including the full pedigree greatly increased both genotype and phase yield and accuracy. The gains were smaller for high coverage individuals compared to low coverage individuals. For example, phase accuracy increased on average from 0.85 to 0.97 for 30x individuals, but increased on average from 0.67 to 0.89 for 1x individuals.

**Figure 2.**
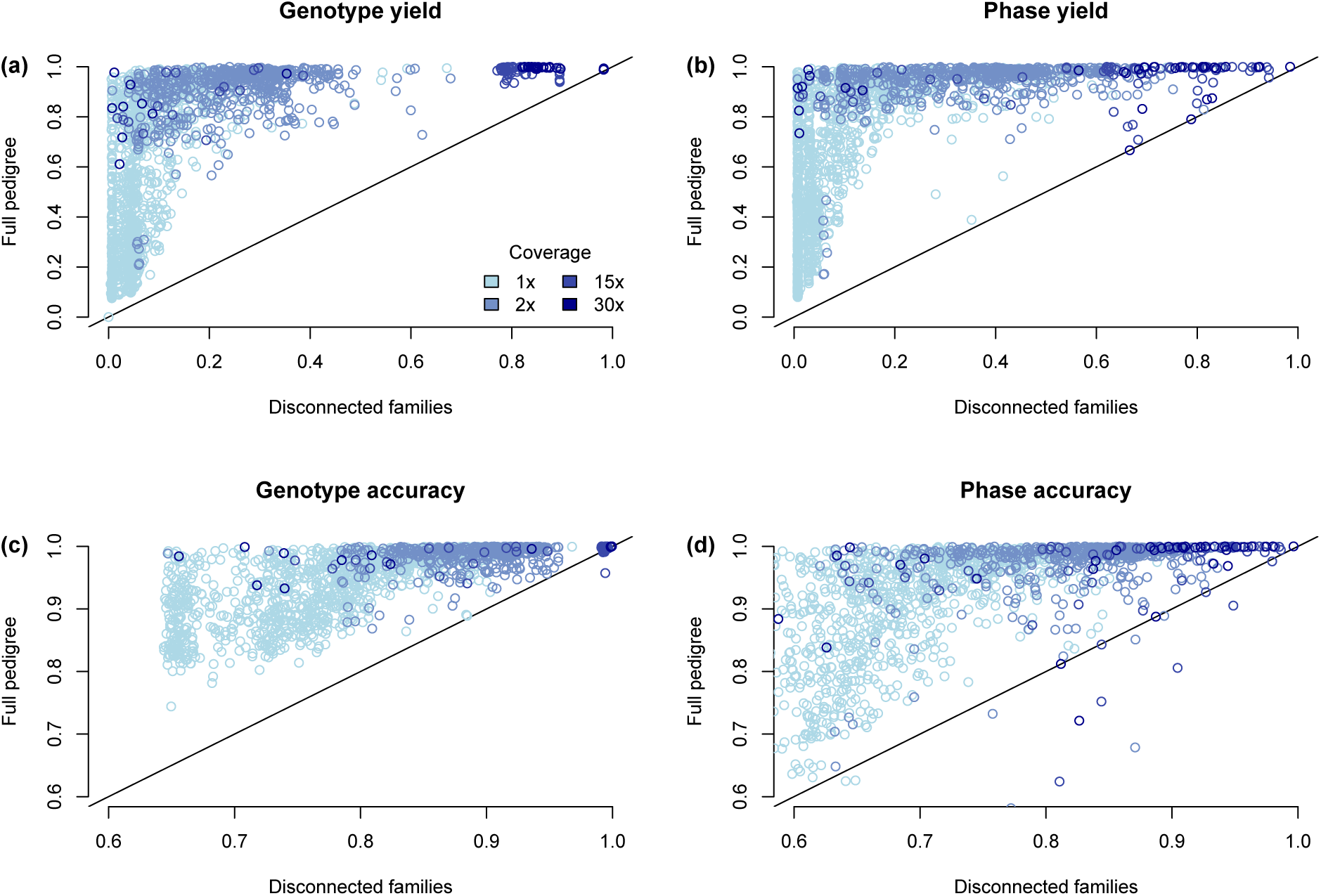
Genotype and phase yield and accuracy when hybrid peeling is run on a series of disconnected families containing a focal individual and its parents and grandparents, or as part of the full pedigree. Panels (a) and (c) give the performance of genotyping individuals, measured either with (a) the genotype yield or (c) the correlation between the true genotypes and the imputed genotype dosages. Panels (b) and (d) give the performance of phasing individuals, measured either with (a) the phase yield, or (c) the correlation between the true phase and the imputed phase.

The gains in accuracy were also not equal for all individuals in the pedigree; some individuals had only a small gain in accuracy, whereas others had a large gain in accuracy. This difference was particularly pronounced for 1x individuals where the phase yield on average increased from 0.11 to 0.67, but the standard deviation increased from 0.13 to 0.28. If all individuals were influenced equally by including the full pedigree, we should expect an increase in mean but not a corresponding increase in standard deviation. The increased variability is a consequence of the different sequencing coverages on relatives who are outside of the immediate family. We found that amount of sequencing coverage on immediate relatives (parents and grandparents) is a good predictor for the phase accuracy of 1x individuals in the disconnected family (r^2^ = 0.37), but is a weak predictor for the phase accuracy of those individuals in the full pedigree (r^2^ = 0.13). In contrast, adding in the sequencing coverage on all ancestors increased our ability to predict accuracy when assessing the phase accuracy in the full pedigree (r^2^ increased from 0.13 to 0.42), compared to when assessing the phase accuracy in the disconnected families, (r^2^ increased from 0.37 to 0.55). The higher overall r^2^ for disconnected families is likely due to the fact that performance in a disconnected family is easier to estimate because of the limited interaction between coverage levels for far away ancestors. A similar pattern of results was found for genotype accuracy and the genotype and phase yields.

### Imputing whole genome sequence

Finally, we analysed the ability of hybrid peeling to impute whole genome sequence data to all non-sequenced individuals in the pedigree. Figure 3 plots the imputation accuracy for every individual as a function of their position in their pedigree. In Table 3 we present the median imputation accuracy stratified by the used sequencing sets and individual’s SNP array genotype status. Overall, we imputed highly accurate allele dosages across the entire pedigree using the *focal plus all low coverage* sequencing set, with an accuracy of 0.987 for individuals with high-density SNP array data, 0.967 for individuals with low-density SNP array data, and 0.881 for non-genotyped individuals. We observed a qualitative difference in imputation accuracy in older individuals. Because of this we stratified results for the first quintile (first 12,919 individuals) and the entire pedigree.

**Figure 3.**
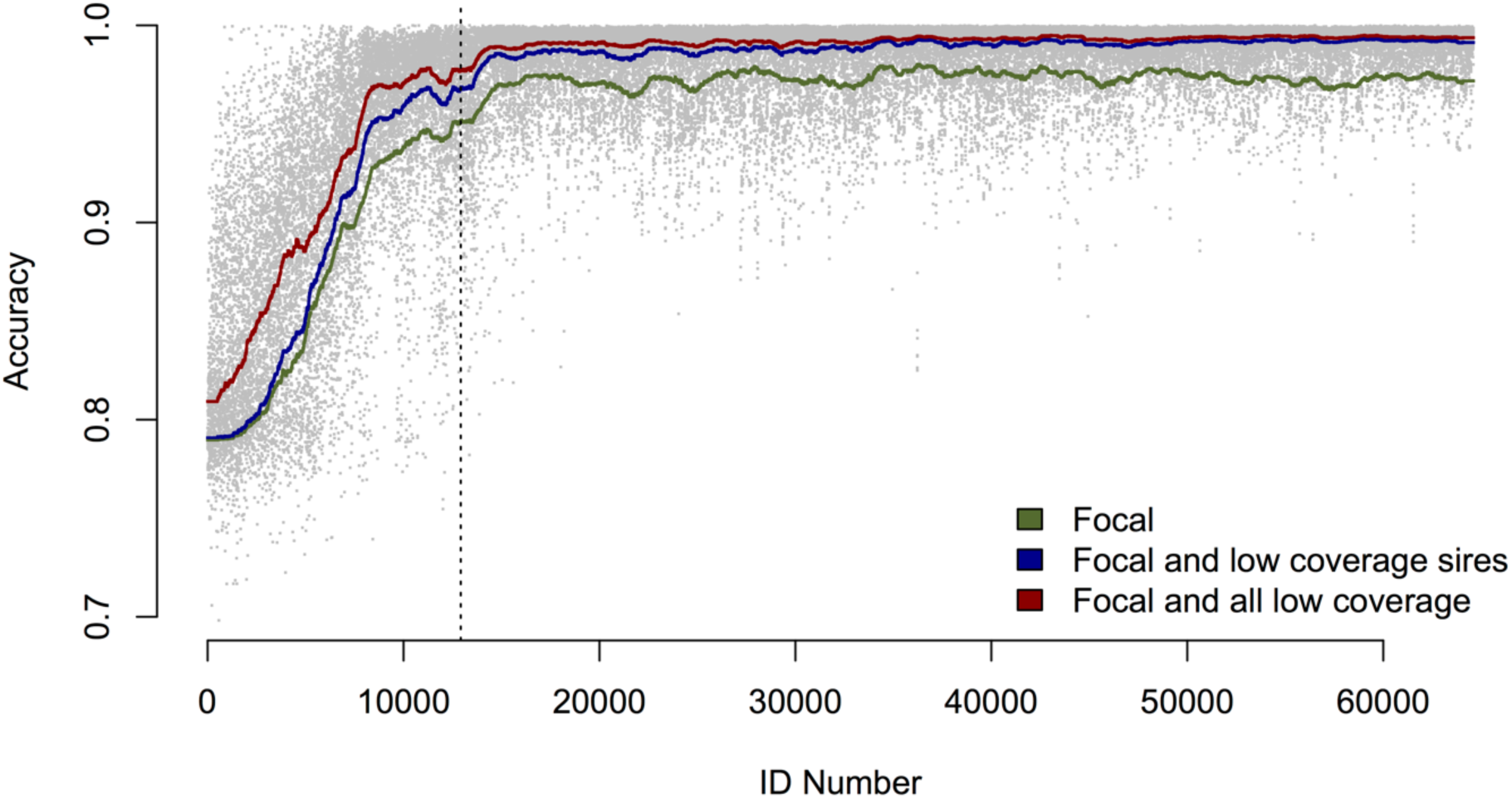
Individual imputation accuracy as a function of birth order (ID number). The green, blue, and red lines track the running average of 1000 individuals when respectively the *focal* individuals, the *focal and low coverage sires*, or the *focal and all low coverage* individuals were used for imputation. The grey dots show results for every individual when the *focal and all low coverage* individuals were used for imputation. The vertical dotted line represents the break between the first quintile of individuals and the remaining four quintiles of individuals.

**Table 3.** Median imputation accuracy for non-sequenced individuals as a function of used sequencing data sets and individual’s SNP array genotype status. These measures were taken over (a) all non-sequenced individuals or (b) the final four quintiles of the population.

| All individuals | High density | Low density | No genotype |
| --- | --- | --- | --- |
| Focal | 0.967 | 0.936 | 0.855 |
| Focal and low coverage sires | 0.983 | 0.952 | 0.863 |
| Focal plus all low coverage | 0.987 | 0.971 | 0.881 |
| Final four quintiles | High density | Low density | No genotype |
| Focal | 0.968 | 0.968 | 0.939 |
| Focal and low coverage sires | 0.984 | 0.985 | 0.953 |
| Focal plus all low coverage | 0.987 | 0.988 | 0.959 |

We observed three trends in imputation accuracy. First, individuals in the first quintile had on average lower imputation accuracy then the rest of the population. When we used the focal plus all low coverage sequencing set the imputation accuracy for the first quintile was 0.908, compared to the average imputation accuracy of 0.970. This decrease in imputation accuracy is due to the lower average sequencing coverage of ancestors for individuals in the first quintile (83x compared to the population average of 308x) and the small number of individuals with high-density SNP array data (0.2% in the first quintile compared to the population average of 70%).

Second, increasing the amount of sequencing resources increased accuracy for all individuals in the population. The largest contribution came from using focal individuals and their parents and grandparents, which gave imputation accuracy of 0.945. Further, adding low coverage sequence data of top sires increased imputation accuracy to 0.963. Finally, adding sequence data of top dams and the remaining low-coverage individuals increased the imputation accuracy only to 0.970, but had a proportionally larger influence on individuals in the first quintile where the imputation accuracy increased from 0.885 to 0.908. The effect is likely due to the fact that 78% of the top dams and top up individuals came from the first quintile.

Third, imputation accuracy for an individual depended on their SNP array genotype status. A comparison of the accuracies depending on their SNP array density is given in Table 3. Overall the difference between having high-density or low-density SNP array data tended to be small, whereas the difference between having SNP array data or not tended to be larger, although this difference decreased in the later generations. For the final four quintiles, the difference between having high-density or low-density SNP array data was negligible (both had an accuracy above 0.987), and the difference between having SNP array data or not was small (0.988 vs 0.959). In comparison, in the first quintile the difference between having high-density or low-density SNP array data was relatively larger (0.983 vs 0.951) and the difference between having SNP array data or not was much larger (0.951 vs 0.868).

### Computational requirements

The computational requirements of hybrid peeling were much less than those for multi-locus peeling. We compared the time it took multi-locus peeling to process the high-density SNP array with 2,000 markers used as an initial step of hybrid peeling to the time it took hybrid peeling to process the remaining sequence with 700,000 segregating loci when using the *focal plus all low coverage* sequencing set. We found that the initial multi-locus peeling step took 823 minutes and 41 GB of memory to process 2,000 SNPs on 64,598 individuals, which translates to 6.3 hours per 1,000 individuals per 1,000 loci. The hybrid peeling step was split across 1000 jobs of 700 SNPs each. Each job took an average of 40 minutes and 2.3 GB of memory, which translates to 53.5 minutes per 1,000 individuals per 1,000 loci and a total of 40,344 minutes (roughly 28 core-days).

## Discussion

In this paper we present a hybrid peeling method for calling, phasing, and imputing sequence data of any coverage in large pedigrees. This method is computationally efficient and enables the benefits of multi-locus peeling to be realised for data sets with tens of thousands of individuals on tens of millions of segregating variants. We evaluated the performance of hybrid peeling in calling and phasing sequence data in a livestock population and in imputing that sequence data to the non-sequenced individuals in the population. Hybrid peeling successfully used the pedigree to propagate information between relatives to call genotypes and phase alleles for individuals with low and high sequencing coverage. Further, calling and phasing these individuals was most effective when the full pedigree was used. Hybrid peeling was also able to whole genome sequence to 60,000 animals with an accuracy above 0.98. We discuss these results in more detail below.

### Hybrid peeling as a genotype calling and phasing method

We found that hybrid peeling effectively used pedigree information to call genotypes and phase alleles in a population of sequenced individuals. When using hybrid peeling, sequence data from an individual’s parents and grandparents increased the number and accuracy of called genotypes and the number and accuracy of phased alleles compared to just using an individual’s own sequence data. We also found that further increases in yield and accuracy could be gained by using more distant relatives. The benefits of using the full pedigree were most apparent for individuals that had low coverage sequencing data (1x and 2x), where in some cases the total genotype yield could rise from 0.1 based on the individuals own sequence data to over 0.9 using the sequence data from the entire pedigree. These results suggest that hybrid peeling could be used to increase the yield of calling and phasing sequence data in pedigrees. The application of hybrid peeling is not limited to individuals with whole genome sequence data, but may also be useful when handling data generated through genotyping via a reduced-representation sequencing (e.g. RAD-seq (Davey et al., 2011) or genotyping-by-sequencing (Elshire et al., 2011; Gorjanc et al., 2015)).

In addition to increasing genotype yield, hybrid peeling also allows for the phasing of many alleles. Using an individual’s own sequence data limits the number of alleles that can be phased to just homozygous loci. In contrast, the number of phased heterozygous loci greatly increased if there was significant sequence coverage on the individual’s parents, grandparents, or even more distant relatives. The ability to accurately phase alleles will be important for downstream imputation and other analyses. Pedigree based methods, like hybrid peeling offer one route for obtaining this information. There are alternative methods that are based on hidden Markov models, e.g. Beagle (Browning and Browning, 2007). These methods phase individual’s alleles by finding shared chromosome segments between an individual and its distant relatives. However, these methods currently do not scale well to performing whole genome sequence phasing and imputation for tens of thousands of individuals (Gilly et al., 2017), making them impractical for many livestock settings.

The power of hybrid peeling comes from its ability to combine sequence data across many related individuals. Hybrid peeling identifies shared chromosome segments between parents and their offspring, and propagates that information to all the individuals who share those segments. In many cases, particularly with low coverage sequence data it is not possible to clearly identify shared chromosome segments. Hybrid peeling handles those cases by marginalizing over the uncertainty of which chromosome was inherited and so potentially increases the accuracy rate over methods that initially require a high accuracy of determination of shared chromosome segments. By marginalizing over uncertainty, hybrid peeling is able to exploit even low coverage sequence data over many generations. When analysing the performance increase between phasing 1x individuals in the case of disconnected families, versus the case of the full pedigree, we found that most reliable indicator of phasing accuracy was the total amount of sequencing coverage for all of the individual’s ancestors, and not the amount of sequencing coverage on the individual’s parents and grandparent, suggesting that hybrid peeling is able to use even distant relatives to phase individuals.

The heavy reliance of pedigree based imputation is both a boon and a curse for hybrid peeling. As we discuss above, using pedigree information can lead to high accuracy, high yield genotype calling and phasing for low coverage individuals. The usefulness of this technique relies on multi-generational pedigree information being available. Although there is some benefit on using sequence information on an individual’s parents and grandparents, the primary benefit comes from aggregating sequencing information across many generations. The availability of multi-generational pedigree information is generally routinely available in commercial livestock populations, but may be less available for human or wild animal populations. When limited pedigree information is unavailable, the performance of hybrid peeling may be less than that of non-pedigree based imputation methods that rely on linkage disequilibrium to call and phase sequence data (VanRaden et al., 2015). There may be some benefit in combining linkage based information with pedigree based information for calling and phasing animals in populations with shallow pedigrees where linkage information between disconnected populations can be exploited. Existing methods have already considered combining linkage based information on the context of multi-locus peeling (Meuwissen and Goddard, 2010), and for using pedigree based information in the context of linkage disequilibrium based calling and phasing algorithms (Chen et al., 2013; O’Connell et al., 2014). Future work is needed to analyse the optimal integration of hybrid peeling with linkage based methods for use in low-depth pedigrees.

### Hybrid peeling as a whole pedigree imputation method

We found that hybrid peeling could effectively use mixed coverage sequence data to impute whole genome sequence into the non-sequenced individuals in the pedigree. For the majority of individuals we obtained imputation accuracy of 0.98. Imputation accuracy was lower at the beginning of the pedigree compared to the end of the pedigree due to the low ancestral sequencing coverage and the high number of individuals genotyped with low-density SNP arrays early in the pedigree. This trend identifies a difficulty that many pedigree based imputation methods face, i.e., it is generally easier to impute children from their parents then it is to impute parents from their children. This difficulty arises from the fact that it is often challenging to phase parents based on their children’s genotype. Doing so requires finding patterns of shared inheritance across multiple offspring, and generally requires many children (Ferdosi et al., 2014). In contrast, it is relatively easy to phase a child’s genotype based on its parents’ genotypes.

One of the more surprising results was the high accuracy observed for non-genotyped individuals. Restricted to the last four quintiles of individuals in the pedigree, non-genotyped individuals had an imputation accuracy of 0.959, which is only slightly less than the 0.988 accuracy for individuals that had high-density SNP array data. The only information that hybrid peeling had for non-genotyped individuals was their position in the pedigree and the list of parents, mates, and offspring. Using this information hybrid peeling was able to accurately reconstruct inheritance of chromosomes across generations, and impute these individuals up to whole genome sequence. The ability of hybrid peeling to impute non-genotyped pedigree members highlights the difference between pedigree and linkage disequilibrium based methods such as Beagle (Browning and Browning, 2007), Impute2 (Howie et al., 2009), or MaCH (Li et al., 2010), which require all individuals to be genotyped at, at least, with a low-density SNP array.

We also noted significant computational gains of hybrid peeling compared to the multi-locus peeling of Meuwissen and Goddard (2010). Both methods scale as O(NL)-linearly with the number of individuals (N) and number of loci (L). However, compared to full multi-locus peeling we found that hybrid peeling ran about 6 times faster and used less memory than full multi-locus peeling. The increased speed stems from not having to update the segregation estimates at each locus. The decreased memory stems from being able to run each locus independently. This means that memory requirements of hybrid peeling scale linearly with the number of individuals O(N), while multi-locus peeling memory requirements scale linearly both with the number of individuals and number of loci O(NL). The gains in speed and memory also lead to practical gains in implementing hybrid peeling. Because each locus is considered independent of the other loci given the segregation estimates, hybrid peeling is trivial to parallelize. Further, the lower memory requirement allows this parallelization to be done on even small machines. Parallelisation meant that although overall imputation time for 700,000 segregating loci on 64,598 individuals took 28 days of CPU time, we were able to run it on a computing cluster in under 24 hours of real time.

## Conclusions

This paper presents hybrid peeling, a computationally tractable multi-locus peeling algorithm for whole genome sequence data. We demonstrated the effectiveness of hybrid peeling in calling, phasing, and imputing whole genome sequence in a large livestock population. We found that hybrid peeling could effectively use multiple generations of variable coverage sequence data to easily increase the yield and accuracy of called genotypes and phased alleles compared to using an individual’s own sequence data. We also found that hybrid peeling could accurately impute whole genome sequence into non-sequenced individuals. We implemented a version of this method in the software package AlphaPeel, which is available from the AlphaGenes website (http://www.alphagenes.roslin.ed.ac.uk). Hybrid peeling has the potential to open the door the routine utilization of whole genome sequence in large pedigreed populations, increasing the accuracy of genomic prediction and the power to detect quantitative trait loci.

## Data availability

Simulated genotype and sequence data are available from the authors upon request.

## Code availability

To perform hybrid peeling we used the software package AlphaPeel, which is available from the AlphaGenes website (http://www.alphagenes.roslin.ed.ac.uk). The code for generating simulated sequence data from genotype data is available from the authors on request.

## Author contributions

AW, GG, and JMH designed the hybrid peeling algorithm. AW and DLW wrote the code. AW and RR designed and ran the simulation study. All authors contributed to writing the manuscript.

